# NG-meta-profiler: fast processing of metagenomes using NGLess, a domain-specific language

**DOI:** 10.1101/367755

**Authors:** Luis Pedro Coelho, Renato Alves, Paulo Monteiro, Jaime Huerta-Cepas, Ana Teresa Freitas, Peer Bork

## Abstract

NGLess is a domain specific language for describing next-generation sequence processing pipelines. It was developed with the goal of enabling user-friendly computational reproducibility.

Using this framework, we developed `NG-meta-profiler`, a fast profiler for metagenomes which performs sequence preprocessing, mapping to bundled databases, filtering of the mapping results, and profiling (taxonomic and functional). It is significantly faster than either MOCAT2 or htseq-count and (as it builds on NGLess) its results are perfectly reproducible. These pipelines can easily be customized and extended with other tools.

NGLess and NG-meta-profiler are open source software (under the liberal MIT licence) and can be downloaded from http://ngless.embl.de or installed through bioconda.

## Introduction

Over the last decade, metagenomics has increasingly been applied for the study of microbial communities. Most work has focused on human-associated habitats^1^, with a particular emphasis on the human gut microbiome^2^,3. However, the same methodologies have been used for studying other host-associated microbiota^4–6^ or the marine microbiome^7^. Due to its size and complexity, several computational approaches have been proposed to handle these data, including bioinformatic pipelines combining different tools and approaches^8–11^.

A typical metagenomics processing workflow can be divided into two distinct phases: in the first phase, raw data is processed (often using prebuilt reference databases) to generate a table of feature abundances (a profile). These features can be either taxonomic or functional annotations. Secondly, these profiles are analysed (often in regards to relevant metadata) using statistical methods and packages such as phyloseq^12^, vegan^13^, or LEfSe^14^. In this work, we are focused on the first phase: namely obtaining functional and taxonomic profiles from raw metagenomic reads.

To this end, we present `NG-meta-profiler`, a collection of pre-configured pipelines based on the domain-specific language NGLess (Next Generation Language for less effortful analysis). Although `NG-meta-profiler` can be used as a standalone tool, the syntax and semantics of NGLess have been designed to be simple and human readable, allowing users to read or create their own pipelines, even without deep bioinformatics and programming knowledge. In other scientific contexts, domain-specific languages have been empirically found to increase productivity and user satisfaction^15,16^. At the same time, NGLess is designed to enable perfect reproducibility of the computational process, an increasingly important concern^17–19^.

In version 1.0^1^, NGLess implements the following tasks: (1) preprocessing, (2) assembly, (3) open-reading frame (ORF) finding, (4) mapping to to sequence databases, (5) filtering of mapping results, (6) profiling (up-to-date taxonomic and functional profiling databases are provided), (7) summary plots.

Using this framework, we have developed `NG-meta-profiler`, a collection of pipelines for taxonomic and functional profiling of metagenomes. These standard analyses can be run with a single command. However, they can also serve as a starting point for customization by the user, including extending them with novel tools.

## Results

### A bioinformatics-aware language leads to faster and more integrated tools

NGLess is a domain-specific language which was designed specifically for next-generation sequence (NGS) processing (see Supplemental File 1 and the online manual for a full description of the language). It contains builtin types which map to concepts in the sequencing domain (e.g., *short read*) as well as primitives that perform common operations (e.g., preprocessing a set of short reads).

The built-in knowledge of the sequence processing domain allows for best-practices to be automatic. For example, our tool always collects quality control statistics without the user having to specify it as an additional computational step.

Domain knowledge enables the interpreter to perform computations more efficiently. For example, even though users write their pipeline script in a purely linear fashion, the interpreter can automatically detect when parallelization opportunities are available and take advantage of multiple processors. When relying on external tools, NGLess can automatically detect possibilities to avoid the use of intermediate files whenever possible, while handling all format conversions internally.

Finally, error detection and reporting are significantly improved by having the tool be semantically aware of its goals. Given that debugging consumes a significant fraction of the time invested in developing computational pipelines, fast error detection can speed up the overall project. For example, it is possible to check whether inputs are readable and outputs are writable prior to starting interpretation. This benefits the user whenever they have made a mistake as errors are detected and reported immediately.

While introducing a novel language implies that the user needs to learn a new tool, the language is designed to be easily understandable to scientists familiar with the field. Alternatively, the Python interface to NGLess allows users familiar with that programming language to access NGLess functionality. Similarly, `NG-meta-profiler` is a command line tool that can be used directly without knowledge of NGLess.

### NG-meta-profiler: a fast metagenomics profiler

NG-meta-profiler is a collection of predefined pipelines for analyzing metagenomic data, resulting in taxonomic and functional profiles. These pipelines are defined in the NGLess language and currently there are workflows available for human, mouse, pig, and dog gut as well as marine metagenomes.

We describe an abridged version of the human gut metagenome profiling pipeline in detail. At a high level, NG-meta-profiler, performs the following operations: (1) preprocess the reads (performing quality-based trimming and filtering of short reads and, if appropriate, discarding reads that align to the host), (2) map the reads against a predefined gene catalog selected for that biome, and (3) use the predefined gene annotations to build a profile (see Fig. 1).

**Figure 1.**
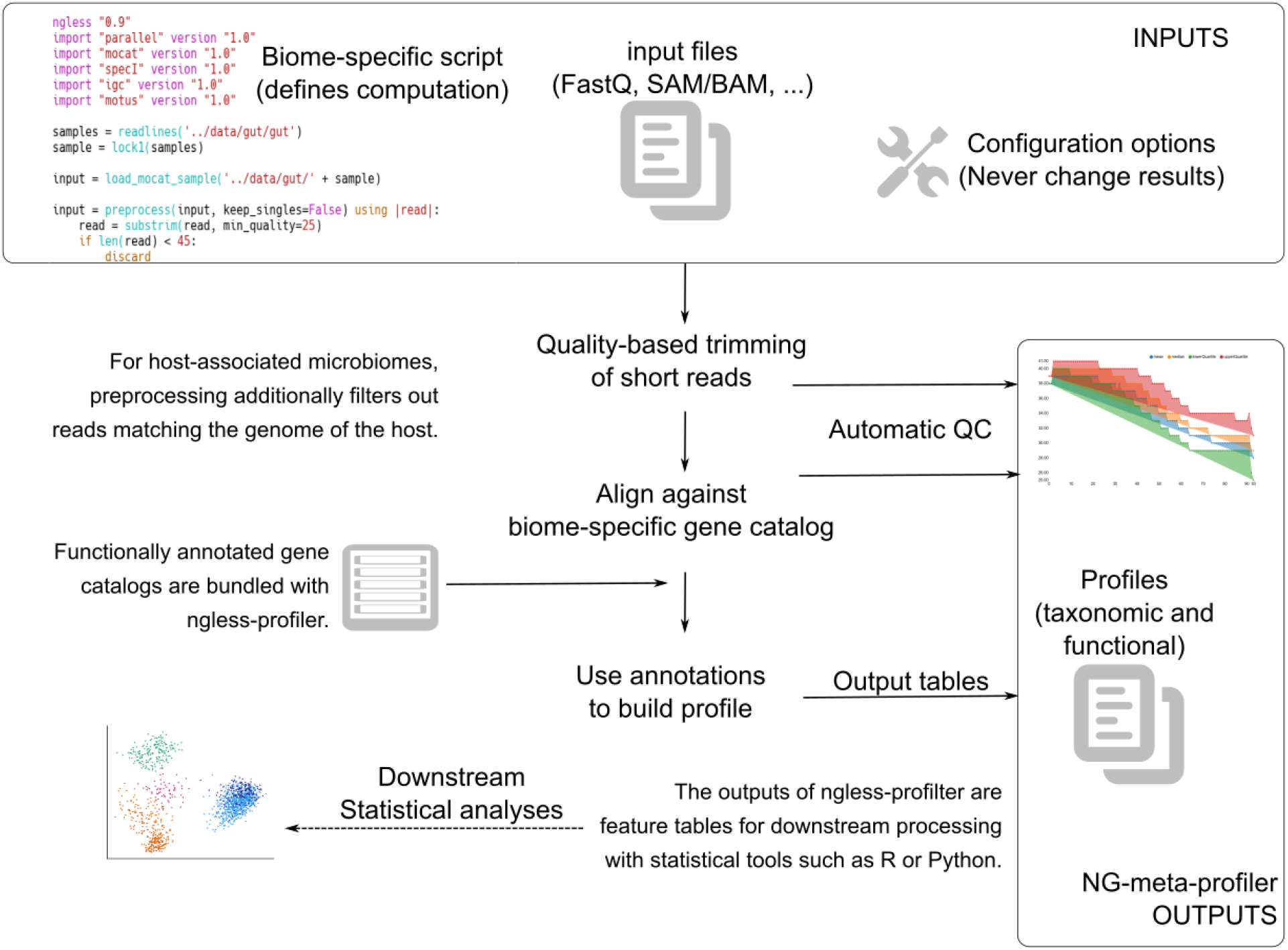
Schematic illustration of NG-meta-profiler.

We aim to describe the operation of the profiler while illustrating the NGLess language. These scripts provide a pre-defined pipeline, but they can also serve as a starting point for customization by the user.

The first element in an NGLess script is the version declaration:

~~~
ngless “0.9”
~~~

The user can then import helper modules. Specifying versions is required for reproducibility. The first module imported is the mocat module. This module allows users to load data that is organized with a sample per directory, an approach that is also used in MOCAT/MOCAT2^8,9^. Secondly, we import the igc module to be able to use the integrated gene catalog (IGC) of the human gut^20^.

~~~
import “mocat” version “0.9”
import “igc” version “0.9”
~~~

The data is loaded using the `load_mocat_sample` function. The input directory is given as the first of the command line arguments (which are stored in an array named ARGV, following the convention from other programming languages):

~~~
input = load_mocat_sample(ARGV[1])
~~~

`preprocess` is a built-in construct to perform quality based trimming and filtering of short reads:

~~~
qc_reads = preprocess(input, keep_singles=False) using |read|:
    read = substrim(read, min_quality=25)
    if len(read) < 45:
        discard
~~~

After preprocessing, it is necessary to reads that match the human genomes. For this operation, we will first align the reads against the hg19 built-in reference:

~~~
human_mapped = map(qc_reads, reference=‘hg19’)
~~~

Spurious alignments (those covering fewer than 45bps or at a lower than 90% nucleotide identity) are now removed, followed by discarding any reads still aligned to the human genome:

~~~
non_human = select(human_mapped) using |mr|:
    mr = mr.filter(min_match_size=45, min_identity_pc=90, action={unmatch})
    if mr.flag({mapped}):
        discard
~~~

The non_human object contains a MappedShortReadSet (the NGLess representation of the information in a SAM file^21^). In order to extract only the sequences (effectively convert it back to a set of FastQ files), the `as_reads` function is used:

~~~
non_human_reads = as_reads(non_human)
~~~

Now, these sequences are mapped to the igc reference (the integrated gene catalogue, imported earlier), and the results are given to count function to obtain a functional profile:

~~~
igc_mapped = map(non_human_reads, reference=‘igc’)
igc_counts = count(igc_mapped,
            features=[‘KEGG_ko’, ‘eggNOG_OG’],
            normalization={scaled})
~~~

Finally, `NG-meta-profiler` saves the result to the file `igc.profile.txt` within the output directory provided by the user:

~~~
write(igc_counts, ofile=ARGV[2] </> ‘igc.profile.txt’)
~~~

### Bundled databases and modules

As part of the first release of `NG-meta-profiler`, we bundle several genomes (including human and mouse) as well as gene catalogs for the human gut microbiome^20^, the marine microbiome^7^, and three non-human mammal microbiomes (pig,^5^, dog^6^ and mouse^4^. These gene catalogs have been functionally annotated with eggnog-mapper^22^ so that functional profiles can be generated by NGLess (see Table 1). In the current version, `NG-meta-profiler` can produce eggNOG orthologous group^23^, KEGG orthologous groups (KO)^24^, SEED^25^, and BiGG^26^ abundance profiles. Additionally, genomes of several organisms (see Supplemental Text 1) are also provided. These are used for filtering out host reads in host-associated metagenomes in `NG-meta-profiler`.

All of these resources are automatically downloaded by NGLess the first time they are used so that the initial download is small (currently 15 MiB) and each user only downloads those resources they effectively use.

**Table 1.** Gene catalogs bundled with NG-meta-profiler.

| Database | Size (million genes) | Comment |
| --- | --- | --- |
| igc | 9.9 | Integrated gene catalog for the human gut<sup>20</sup> |
| om-rgc | 40 | Ocean microbial gene catalog<sup>7</sup> |
| mouse-gut | 2.6 | Gene catalog of the mouse gut<sup>4</sup> |
| pig-gut | 7.7 | Gene catalog of the pig gut<sup>4</sup> |
| dog-gut | 1.2 | Gene catalog of the dog gut<sup>6</sup> |

Several operations in NGLess are performed with bundled software. *De novo* assembly is performed using MEGAHIT^27^, which has been found to perform well for metagenomics^28,29^. Open reading frame (ORF) finding is performed with Prodigal^30^. By default, mapping is performed using bwa^31^, but minimap2^32^ is also available. Additional built-in modules provide extra functionality as shown in Table 2.

**Table 2.**
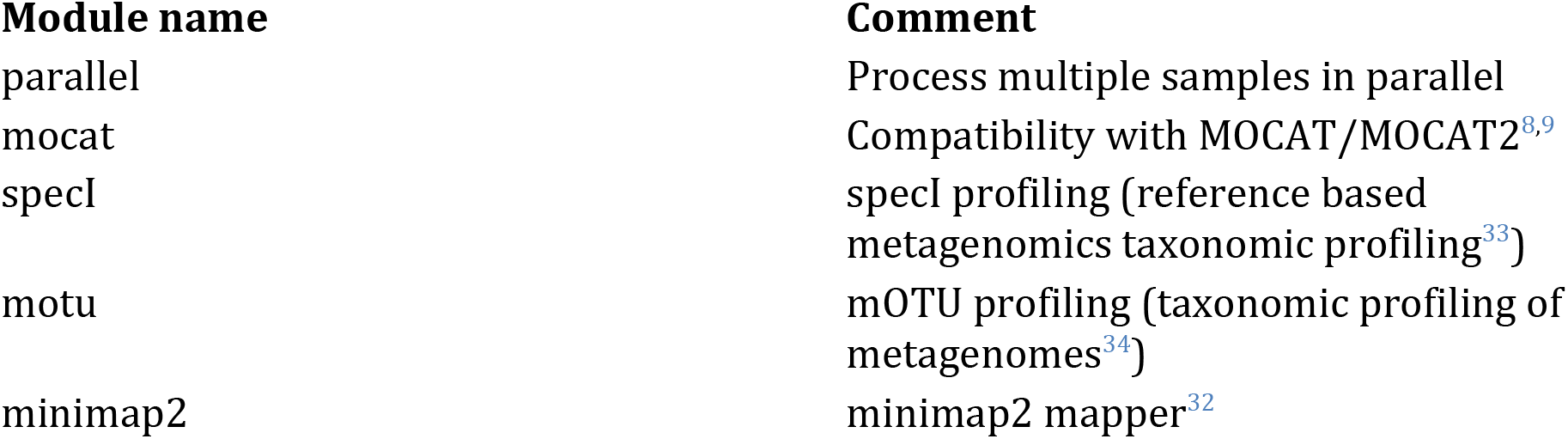
NGLess builtin modules that add extra functionality.

### Benchmarking

We compared the performance of `NG-meta-profiler` with both MOCAT2^9^ and a pipeline based on calling bwa without preprocessing data and htseq-count^35^ for profiling human gut^36^ and ocean metagenomes^7^. In all cases, we used the `NG-meta-profiler` databases and set parameters so that the results are identical, up to rounding errors. For this benchmark, 8 threads were used (except for the htseq-count software which only supports a single thread).

Results are presented in Fig. 2 (the Supplemental Software contains a preprocessed table with the full results). `NG-meta-profiler` clearly outperforms the other solutions in this task. When compared to MOCAT2, the `NG-meta-profiler` runs 11x faster in the marine benchmark, and 2.6x in the human gut metagenome benchmark.

**Figure 2.**
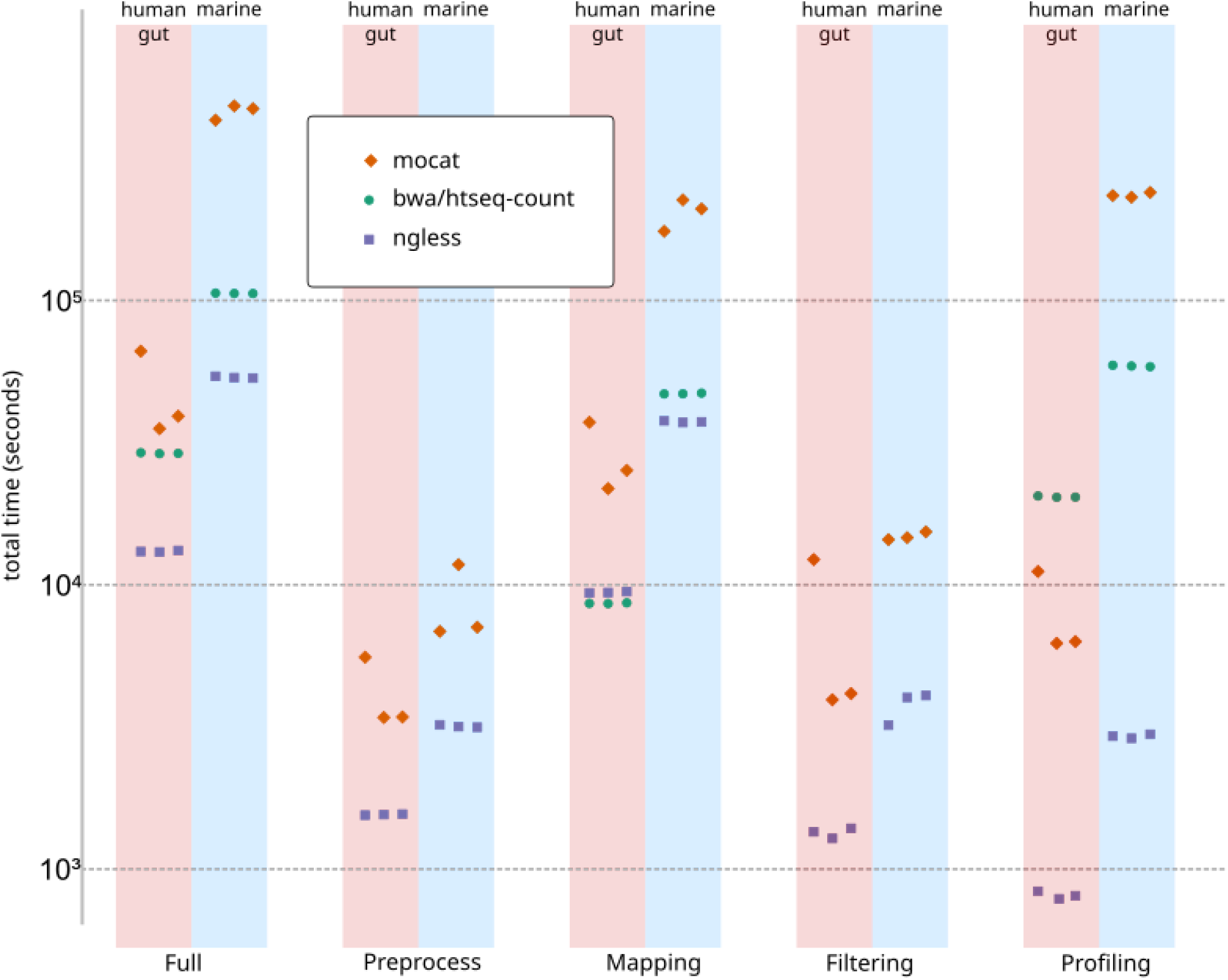
Timing comparison of NGLess and other tools. Three replicates are shown for each tool. The bwa/htseq-count pipeline does not include Preprocess and Filtering steps.

The larger ratio in the larger ocean microbial gene catalog (40 million genes) compared to the integrated gene catalog used for profiling the human gut (10 million genes) is evidence that NGLess scales better to very large catalogs.

Note that running the full `NG-meta-profiler` pipeline takes less time than the sum of the individual steps as the pipeline is optimized as a whole (e.g., it avoids generating unnecessary intermediate files).

### Pipelines developed with NGLess are reproducible

Unlike tools which are based on traditional programming languages, NGLess is designed from the ground up with reproducibility as a goal.

As seen above, every pipeline defined with NGLess includes a version declaration and every imported module specifies the particular version which is being imported. Documenting the version of all the tools and dependencies used for a given analysis is considered a best practice^37,38^, but is not always followed. By making it a requirement within the script, NGLess ensures that this best practice is adopted. At the same time, NGLess lowers the necessary effort when compared to having to record the version of all tools and dependencies manually. For example, the versions of samtools^21^ and bwa^31^ used internally (and shipped with NGLess) are implicitly fixed by specifying the NGLess version. To encourage that credit be given to the original authors, NGLess will print out references for any tools that are used, asking the user to cite them in publications.

Furthermore, although external configuration and command line options may change *how* results are computed (e.g., how many threads to use, where to store temporary files), the results do not depend on any information outside the script. This separation of implementation details from the data processing specification has the added potential of making the resulting code easier to port between systems^39^.

### Extensibility and integration into the wider ecosystem of bioinformatics tools

Pipelines defined with NGLess are easily extensible. We encourage users of `NG-meta-profiler` to customize these pipelines to their specific problems and to extend them as desired. Functions can be added to the language based on external software by specifying the interface in a text format and importing it from the main script.

In May 2018, we opened up to the community a repository of external modules (https://github.com/ngless-toolkit/ngless-contrib) where contributions of new integrations are accepted. At the moment, integration of MetaPhlAn2 (a tool for profiling microbial species based on taxa-specific marker genes^40^) and salmon (a tool which generates abundance profiles based on k-mers^41^) are available.

Alternatively, NGLess based analyses can be integrated into larger pipelines. Most existing bioinformatics pipelines are performed with command-line based software. To facilitate integration with existing tools, we provide several tools based on NGLess with a command line interface. In addition, we provide Common Workflow Language (CWL) descriptions of these tools which enable their use as part of CWL workflows^42^.

NGLess scripts can (with some limitations) be automatically exported as CWL tools in order to be embedded in larger projects. When passed the `--export-cwl` option, NGLess will output a CWL description of a given script which can then be embedded in a larger CWL-based pipeline.

Finally, NGLess can also be used as an embedded language within Python, a programming language that is widely used for scientific computing^15^. With this interface, NGLess-based pipelines can be defined by Python-based scripts.

## Discussion

NGLess puts forward a different approach for defining data analysis pipelines: the use of a domain-specific language for sequence analysis. Several tools had already used a domain-specific language to define a computational pipeline. The classical Make tool was originally designed for compiling software, but has been used as the basis of a scientific pipeline system^43^ uses its own internal language, as do the more modern pipelining tool Snakemake^44^ and nextflow^45^. These tools operate by organizing the computation around calls to command-line software. As such, they are fully generic and can be used in a wide range of problems. NGLess trades off this generality to achieve higher usability within its problem domain.

Using this framework, we developed `NG-meta-profiler` which generates taxonomic and functional profiles from metagenomes based on prebuilt gene catalogs that are provided with the tool. When compared with other alternatives, `NG-meta-profiler` performs reference-based functional and taxonomic profiling much faster. Furthermore, this collection of scripts can be easily adapted and extended by the users within the NGLess framework to perform novel functions.

## Software and data availability

NGLess is available as open source software at http://github.com/ngless-toolkit/ngless. Additionally, NGLess is available as a bioconda package^46^ and in container form (through biocontainers^47^). Documentation and tutorials can be found at http://ngless.embl.de. All the databases mentioned can be downloaded automatically with the NGLess tool or accessed using the DOI 10.5281/zenodo.1299267.

Jug^48^ based scripts to download the data and run the benchmarks are available at https://github.com/ngless-toolkit/NGLess2018benchmark. This repository also contains the results of running the benchmarks on our servers and all downstream processing. The benchmarks were run on an Intel Xeon (CPU model “E7- 4830 @ 2.13GHz”) with 32 cores (64 virtual cores).

The sequence datasets used for the benchmark are available from the European Nucleotide Archive (ENA) (accession numbers: SAMEA2621229, SAMEA2621155, SAMEA2621033, SAMEA2467039, SAMEA2466896, and SAMEA2466965).

## Acknowledgements

The authors thank João Carriço (Instituto de Medicina Molecular, University of Lisbon) as well as the members of the Bork lab for helpful comments. We thank Anna Głazek (EMBL) for several patches to the code. Beta users of NGLess are thanked for their feedback and bug reports.

Funding was provided by the European Union’s Horizon 2020 Research and Innovation Programme (grant #686070; DD-DeCaF), the European Research Council (ERC) MicrobioS (ERC-AdG-669830), the Fundação para a Ciência e a Tecnologia (grant EXCL/EEI-ESS/0257/2012; DataStorm: Large-Scale Data management in cloud environments), and the European Molecular Biology Laboratory (EMBL).

## Author contribution

LPC conceived of the project and wrote the first draft of the manuscript. LPC, ATF, and PB coordinated and oversaw the study. LPC, RA, JHC, and PM developed the software and associated databases. LPC, RA, PM, JHC, and PB drafted the manuscript and supplemental text. LPC and RA designed and performed the benchmark study. All authors contributed to the review of the manuscript before submission for publication. All authors read and approved the final manuscript.

## Footnotes

1 The currently available version is named 0.9. We will to rename it to version 1.0 upon acceptance of this manuscript for publication.

